# The tricellular junctional protein Ildr2 helps maintain glomerular podocyte architecture

**DOI:** 10.1101/2024.05.03.592393

**Authors:** Gary F. Gerlach, Yubin Xiong, Lori L. O’Brien

## Abstract

Podocytes are highly specialized epithelial cells of the glomerular filtration barrier essential for maintaining proper kidney function. Their intricate cellular projections, called foot processes, wrap around the capillary endothelium, interdigitate, and connect to one another via slit diaphragm intercellular junctions to form a sieve-like barrier. Compromise to this unique podocyte architecture can lead to altered glomerular function and eventual kidney disease. We previously identified Immunoglobulin-like domain containing receptor 2 (Ildr2) as a component of the podocyte foot process through proteomic analyses. Ildr2 is a tricellular tight junction constituent and localizes to distinct puncta in podocytes that likely represent these specialized junctions. However, the significance of tricellular tight junctions and Ildr2 to podocyte integrity is unknown. To this end, we generated a conditional knockout mouse with podocyte-specific deletion of *Ildr2* (*Podocin-Cre^tg/+^;Ildr2^fl/fl^*or *Ildr2^PodKO^*). Kidneys from *Ildr2^PodKO^* adult animals show disruptions to podocyte foot process architecture including effacement and the unique formation of electron dense strands connecting subsets of processes. Additionally, the glomerular basement membrane was significantly thicker in conditional knockouts and histological analyses revealed glomerular collagen and glycoprotein accumulation. Surprisingly, *Ildr2^PodKO^* animals displayed no significant albuminuria. When *Ildr2^PodKO^* mice were challenged with a stress to kidney function, immunohistochemistry revealed podocyte loss from a subset of glomeruli with concomitant detection of podocytes in urine, although function was still preserved. Collectively, our data provide novel insights into the function of Ildr2 and suggest tricellular junctions help preserve podocyte architecture but are likely not necessary to maintain proper filtration in no or low stress physiological states.

## Introduction

To filter blood and enable the production of urine, the kidney relies on the filtering component of the nephron, the glomerulus. The glomerular filtration unit consists of fenestrated endothelial capillary loops which are covered by podocytes, in addition to the intervening glomerular basement membrane (GBM). Podocytes are architecturally unique epithelial cells that play a crucial role in maintaining integrity and function of the glomerular filtration barrier. Extending from the podocyte cell body are numerous protrusions termed foot processes. Foot processes wrap around the capillaries and interdigitate, with a protein bridge forming between adjacent processes. This protein bridge, or slit diaphragm, is a bicellular junction which contributes to the formation of the glomerular filtration barrier by establishing a physical sieve between adjacent podocytes and their foot processes^1–3^. These junctions enable the selective filtration of plasma components and maintain podocyte architecture^4,5^. Disruption of slit diaphragms and thereby foot process integrity can lead to foot process effacement, impaired barrier function, and loss of cell adhesion, all hallmarks of podocytopathies which make up a significant proportion of kidney disease cases^6,7^. Therefore, the nature of the podocytes’ function makes their foot processes and specialized junctions vital to maintaining human health.

Recent advances in microscopy and image reconstruction have brought to light new observations on foot process architecture^8–11^. Each podocyte displays an extensive arborized foot process network which interdigitates distinctly with another podocyte. To acquire this unique organization, junctional rearrangements occur during podocyte development as these cellular protrusions grow and the slit diaphragm is established^9^. Observations associated with mature podocyte morphology such as a ridge-like prominence and “filamentous cell-cell contacts” highlights areas where additional junctional components may reside outside the slit diaphragm to help maintain integrity^9–11^. Tricellular tight junctions (tTJs) are specialized intercellular connections that occur at the convergence of three neighboring cells^12^. They are important for barrier function and have been implicated in diseases such as familial deafness^12,13^. Tricellulin and angulins are the two major proteins known to localize to tTJs^12^. Disrupting components of tTJs can lead to redistribution of tTJ proteins to bicellular TJs^14^. While tTJs have been studied in other epithelial cells, their presence and significance in podocytes has not been characterized, although there is evidence for their presence during podocyte maturation as junctions are rearranged^9^. Given the critical role of podocytes in maintaining glomerular filtration barrier function, it is plausible that tTJs may contribute to the structural integrity and permeability properties of the filtration barrier.

We previously created a novel mouse model to generate a spatially localized proteome for the podocyte foot process^15^. Ildr2 was identified as a top hit in our mass spectrometry analyses. Ildr2 is a transmembrane protein that belongs to the immunoglobulin superfamily and has been implicated in various cellular processes, including splicing, transcriptional regulation, cell adhesion, and immune modulation^13,16–18^. Its extracellular region contains immunoglobulin-like domains that likely facilitate binding to other cell surface molecules, promoting the formation and maintenance of cell-cell contacts^13,16^. Within the slit diaphragm, the immunoglobulin domains of Nephrin and Neph1 (Kirrel) interact and form a mesh-like organization which maintains a stable podocyte architecture and junctional barrier integrity^19,20^. Ildr2 is a member of the angulin family along with the related protein Ildr1 and localizes to tTJs of epithelial cells^13^. In podocytes, Ildr2/ILDR2 localizes to punctate foci, which partially overlap with podocin, and likely represent sites of tTJs^15,21^. Ildr2/ILDR2 displays altered localization dynamics during development, aging, and disease, in addition to accumulation of the protein in diseased and aged kidney samples, suggesting altered junctional rearrangements are associated with changes in podocyte architecture^15,21^. Intriguingly, Ildr1 is localized to tTJs of nephron distal tubules and knockout studies have revealed a role for these tTJs in paracellular water permeation. The *Ildr1* knockout mice have urine concentrating defects due the significant increase in water permeability, highlighting a critical role for tTJs in the nephron^22^.

Given the importance of cell-cell junctions such as the slit diaphragm in maintaining the integrity of the glomerular filtration barrier, and the importance of tTJs in other cells of the kidney, we generated a podocyte-specific knockout of *Ildr2* (*Podocin-Cre^tg/+^;Ildr2^fl/fl^* or *Ildr2^PodKO^*) to investigate its functions^23,24^. We identify disruptions to podocyte architecture and histological changes but surprisingly kidney function is preserved in these animals. *Ildr2^PodKO^* podocytes are slightly more prone to detachment with increased stress on kidney function although barrier function remains intact. Our findings highlight the importance of Ildr2 and tTJs to maintaining podocyte architecture but suggest Ildr2 is not necessary to maintain the glomerular filtration barrier.

## Results

### Ildr2 deletion from podocytes results in ultrastructural changes to the filtration barrier

To generate a podocyte specific knockout of *Ildr2*, we utilized the *Podocin-Cre^tg^* line crossed to *Ildr2^flox^*mice to generate controls (*Podocin-Cre^tg/+^;Ildr2^fl/+^*or *Ildr2^Pod/+^*) and conditional knockout animals (*Podocin-Cre^tg/+^;Ildr2^fl/fl^* or *Ildr2^PodKO^*)^23,24^. Our controls included the *Podocin-Cre* transgene to account for any effects of the transgene insertion. To assess the effects of *Ildr2* loss on podocyte architecture, we utilized transmission electron microscopy (TEM) to investigate ultrastructural changes. In *Ildr2^PodKO^* animals we observed interspersed areas of significant foot process effacement (Fig. 1A, arrow, right panel) as opposed to the normal organization of control animals (Fig. 1A, left panel). The glomerular basement membrane (GBM) was also thickened compared to controls and occurred in areas both with effacement and normal foot process organization (Fig. 1B, brackets). Quantitation confirmed a significant difference in GBM thickness (Fig. 1C). Despite these ultrastructural changes, slit diaphragms were still largely present in *Ildr2^PodKO^* animals (Fig. 1B,D). Notably though, *Ildr2^PodKO^* podocytes displayed foot processes with electron dense filaments emanating from the entire surface (Fig. 1D, arrows), seemingly connecting to adjacent and overlying foot processes and differing from the electron dense slit diaphragm (Fig. 1D, arrowhead).

**Figure 1.**
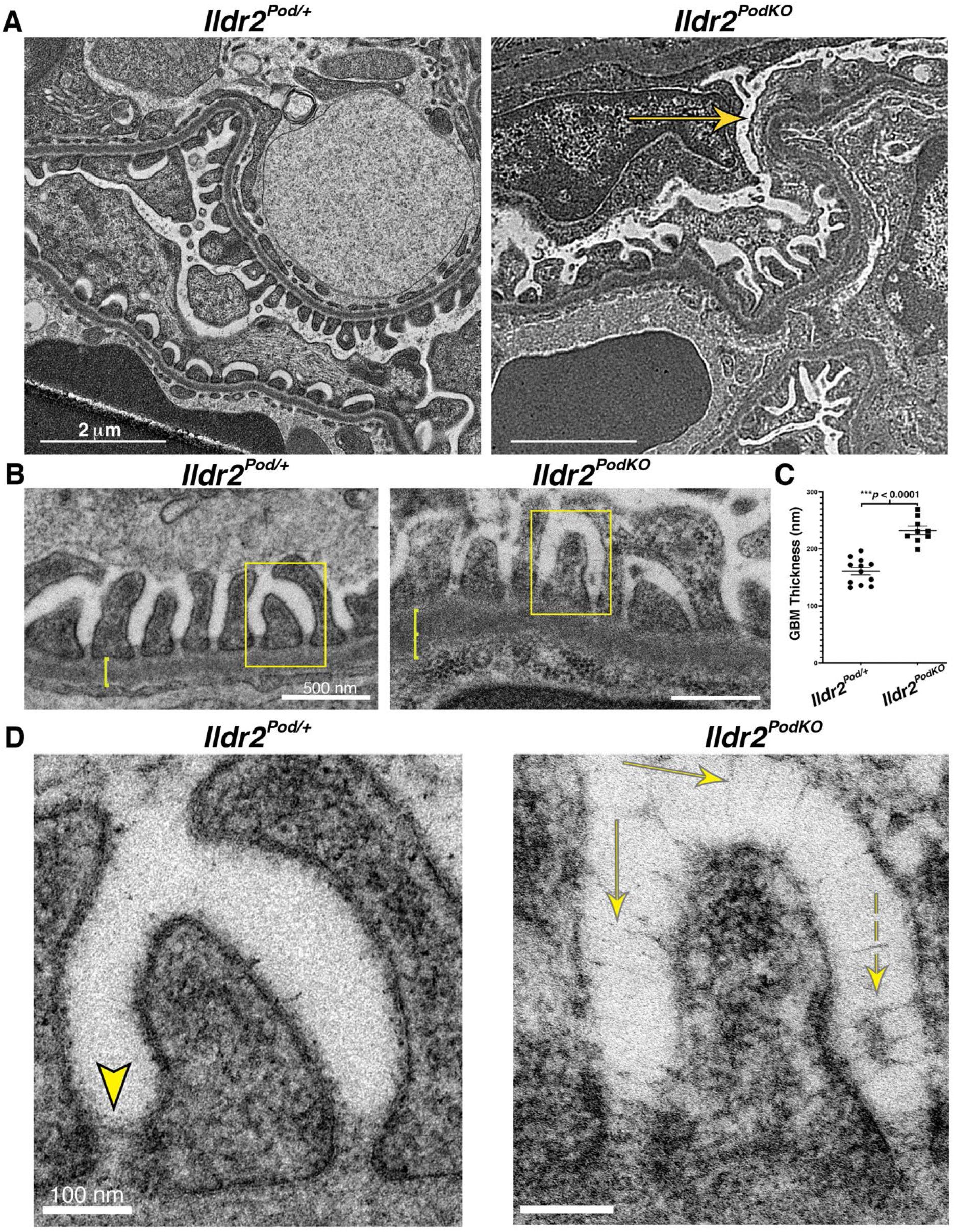
*Ildr2* deletion from podocytes results in ultrastructural changes to podocyte architecture. A) TEM of control *Ildr2^Pod/+^* podocytes show normally spaced foot processes atop the GBM (left panel). In contrast, *Ildr2^PodKO^*kidneys show foot processes effacement (yellow arrow, right panel), Scale bar=2μm. B) *Ildr2^Pod/+^* kidneys show normal GBM tri-layered morphology (lamina rara externa layer, lamina densa, and lamina rara interna layer) atop the endothelium (left panel)*. Ildr2^PodKO^* kidneys show less well-defined boundaries between the GBM layers and a thickening of the GBM (yellow brackets). Scale bar=500nm. C) Quantification of GBM longitudinal thickness shows an increase in *Ildr2^PodKO^* kidneys (mean thickness=230nm±10nm) compared to *Ildr2^Pod/+^* kidneys (mean thickness=160nm±10nm). D) Enlarged yellow boxes from (B) highlighting filtration slit architecture. Control animals (left panel) show a slit diaphragm running roughly parallel to the GBM, between two adjoining foot processes (yellow arrowhead). In contrast *Ildr2^PodKo^* mice (right panel) have numerous protrusions extending from a single foot process at varying angles, in some cases spanning the filtration space (yellow arrows). Electron dense areas associated with the strands are also observed (dotted yellow arrow). Scale bar=100nm.

### Ildr2^PodKO^ kidneys show histological changes to the glomerulus

Histological analyses were performed to identify whether the ultrastructural defects in podocyte architecture and thickening of the glomerular basement membrane resulted in detectable phenotypes such as increased collagen deposition. Immunohistochemical analysis of collagen IV revealed a qualitatively minor increase in collagen deposition within *Ildr2^PodKO^* glomeruli at 3 months of age (Fig. 2A, arrow). Sirius Red staining of histological sections from 7-month-old mice validated the increase in collagen within glomeruli which was significantly increased in *Ildr2^PodKO^* mice compared to controls (Fig. 2B,C). Periodic Acid Shift (PAS) staining of these 7-month samples highlighted thickening around Bowman’s capsule and the urinary pole of *Ildr2^PodKO^* glomeruli (Fig. 2D) suggesting potential collagen deposition as PAS detects proteoglycans, glycolipids, and mucins which are enriched in basement membranes. Surprisingly, despite these ultrastructural and histological changes, we did not detect any significant increase in proteinuria within any cohort of *Ildr2^PodKO^*animals compared to *Ildr2^Pod/+^* controls (Fig. S1).

**Figure 2.**
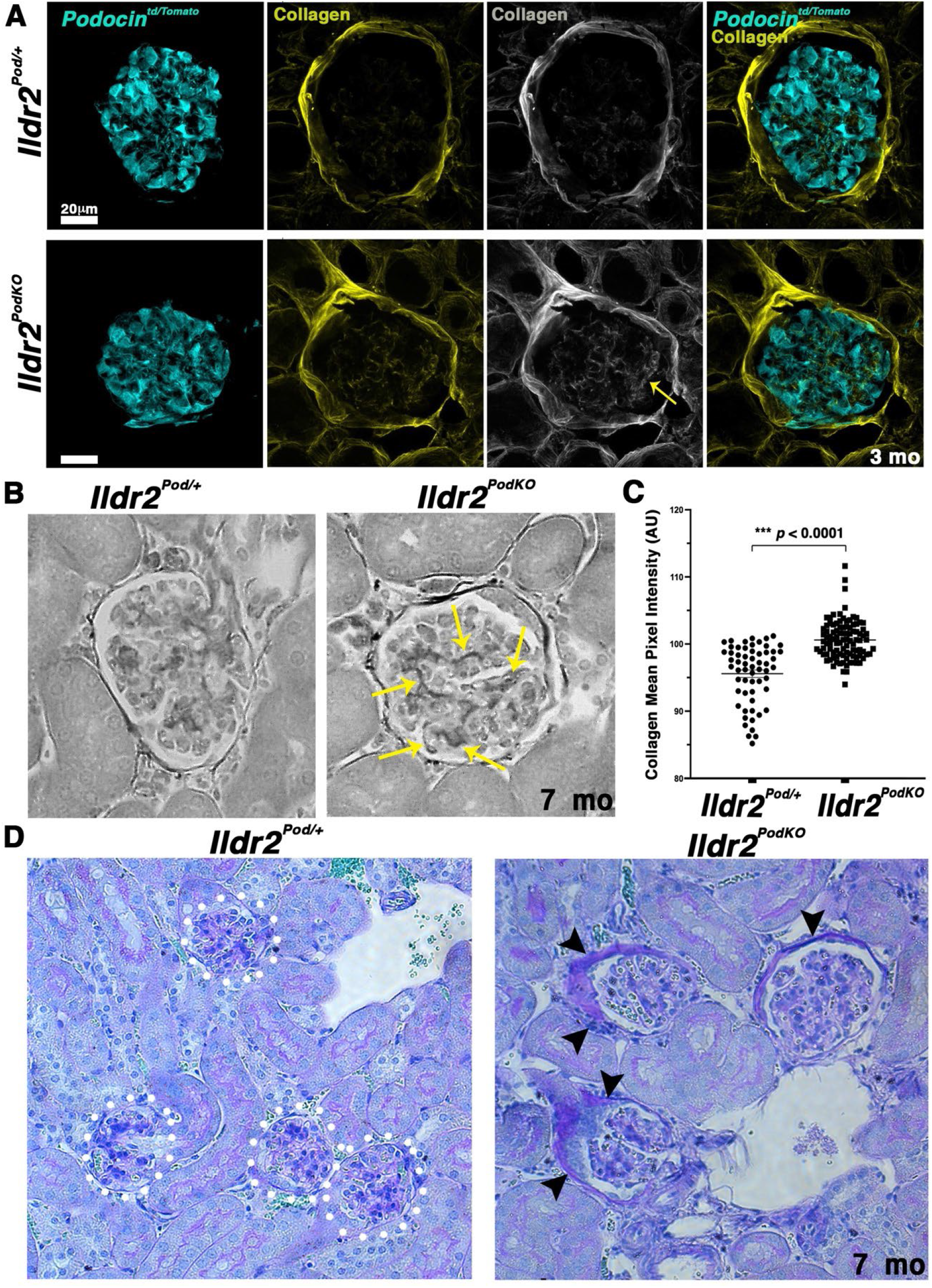
*Ildr2^PodKO^* kidneys show histological glomerular abnormalities. A) Immunohistological analysis of Collagen IV shows a small qualitative increase in collagen within *Ildr2^PodKO^*glomeruli compared to controls at 3 months. Podocytes are labeled with the tdTomato reporter crossed onto the *Ildr2^PodKO^* and *Ildr2^Pod/+^* backgrounds and is driven by *Podocin-Cre^tg^* (*Podocin^tdTomato^*). Collagen IV (yellow/grey) and Tomato+ podocytes (cyan) are labeled. Scale bar=20 µm. B) Histological analysis with Sirius Red (rendered in greyscale) shows increased collagen within glomeruli of 7-month-old *Ildr2^PodKO^*mice (yellow arrows). C) Quantification of mean pixel intensity of glomerular Sirius Red histology in control *Ildr2^Pod/+^* (n=59 glomeruli from 3 mice) and *Ildr2^PodKO^* (n=99 glomeruli from 4 mice) kidneys. D) Periodic acid– Schiff (PAS) histology shows increased signal around Bowman’s capsule and the urinary poles of 7-month-old *lldr2^PodKO^* animals (arrowheads). Control glomeruli are outlined by dotted white circles.

### The localization of junctional components is unaltered in Ildr2^PodKO^ glomeruli

As junctional rearrangements can occur with podocyte injury and disease, we examined whether changes in distribution or levels of junctional proteins occur in our *Ildr2^PodKO^* animals^4,6,25,26^. Immunohistological analyses revealed normal localization and qualitatively similar levels of slit diaphragm components podocin and nephrin (Fig. 3, left panels). We next examined the distribution of ZO-1, a component of bicellular tight junctions important for maintaining the filtration barrier^27^. Like the slit diaphragm proteins, ZO-1 localization and levels were similar between *Ildr2^PodKO^*glomeruli and *Ildr2^Pod/+^* controls (Fig. 3, right panels). Additionally, qPCR revealed no significant differences in the expression of either *Nphs2* (podocin) of *Tjp1* (ZO-1) (Fig. S2). Tricellulin (Marveld2) is a known component of tTJs^12,13^, however, we were unable to identify a working antibody to assess tTJ components. Therefore, we interrogated the localization of vinculin, a cytoskeletal protein involved in cell-cell adhesions, which has well established antibodies. Vinculin is enriched at tTJs to help maintain their integrity and also plays an important role in maintaining the glomerular filtration barrier^12,28–30^. However, we did not observe any significant differences in vinculin distribution or amounts between *Ildr2^PodKO^* kidneys and controls (Fig. 3, right panels). Collectively, cellular junctions appear to be preserved despite *Ildr2* loss.

**Figure 3.**
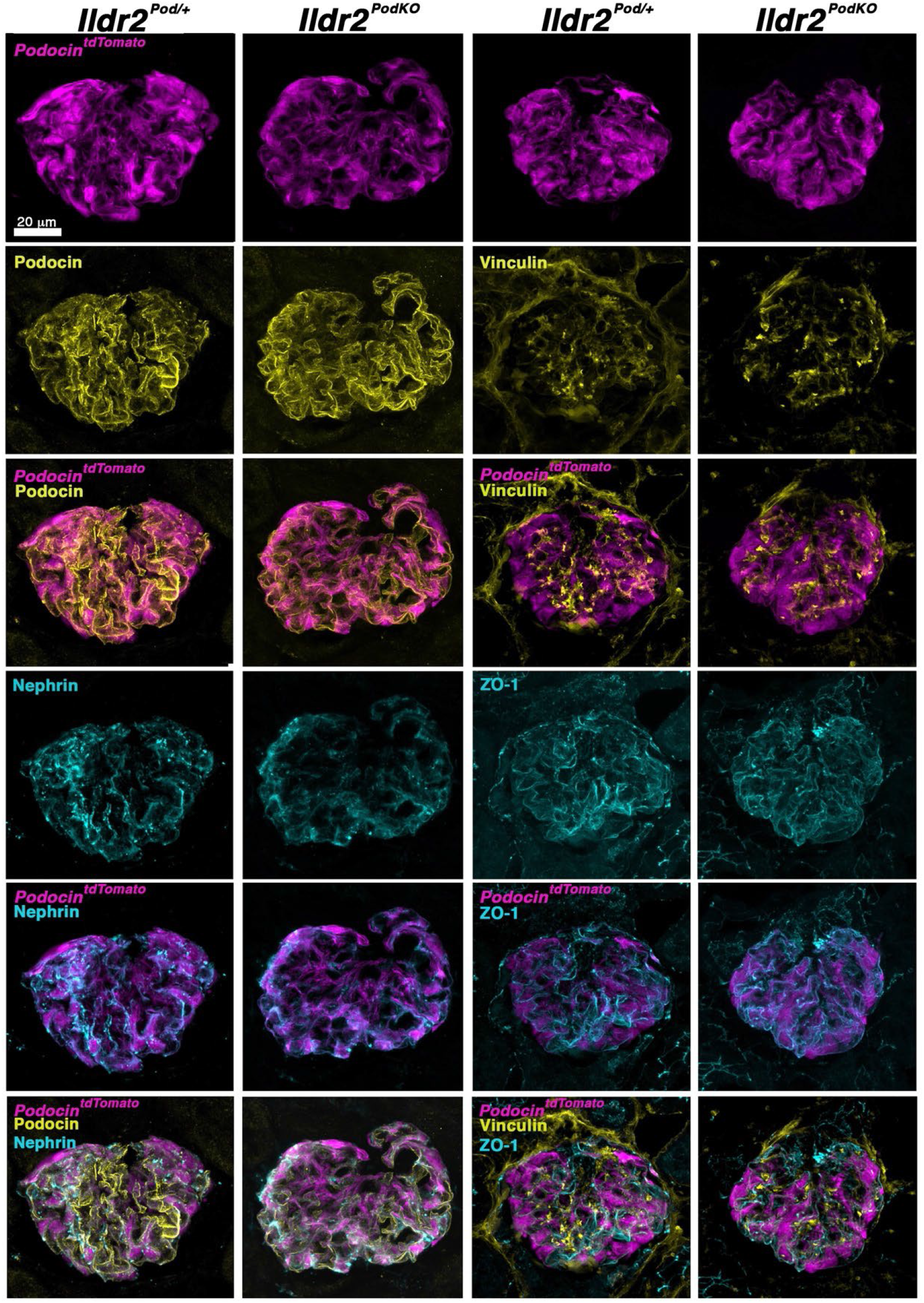
Loss of *Ildr2* does not disrupt the slit diaphragm or other junctional components. Immunohistological analyses of podocin, nephrin, vinculin, and ZO-1 from *Ildr2^PodKO^* and *Ildr2^Pod/+^* kidneys does not identify any significant differences in protein localization or levels. Podocytes are labeled with the tdTomato reporter crossed onto the *Ildr2^PodKO^* and *Ildr2^Pod/+^* backgrounds and is driven by *Podocin-Cre^tg^* (*Podocin^tdTomato^*). Podocin (yellow), Tomato+ podocytes (magenta), and nephrin (cyan) are labeled in the two left panels. Vinculin (yellow), Tomato+ podocytes (magenta), and ZO-1 (cyan) are labeled in the two right panels. Scale bar=20 µm.

### Challenge to kidney function induces podocyte loss in Ildr2^PodKO^ animals

With the absence of proteinuria in *Ildr2^PodKO^* animals, as well as their relative preservation of junctions despite ultrastructural changes to the podocytes, we wondered whether *Ildr2^PodKO^* podocytes may be sensitized. Therefore, we set out to challenge the animals by forcing the kidney to function at high capacity through injection of a 10% of body weight saline solution. The animals were analyzed at 24 hours post-injection to determine if this challenge was sufficient to induce phenotypic and/or physiological changes. The overall volume of urine collected was similar between *Ildr2^PodKO^* and *Ildr2^Pod/+^* animals (Fig. S3A). Intriguingly, immunohistological assessment of the kidneys revealed a portion of *Ildr2^PodKO^* glomeruli that appeared to be lacking podocytes, often retaining only small clusters of cells within the Bowman’s space (Fig. 4A). Quantitation of the abnormal glomerular phenotypes revealed a significant increase (13% for *Ildr2^PodKO^* vs. 1% for *Ildr2^Pod/+^*) in the percentage of glomeruli with missing podocytes (Fig. 4B). To determine whether the challenge was leading to podocyte detachment, we examined collected urine by Western blot for the presence of podocyte proteins. Wt1 and podocin, as well as the Tomato lineage marker crossed onto the *Ildr2^PodKO^* background, were identified in the urine of a subset of *Ildr2^PodKO^* animals (Fig. 4C, Fig. S3B,C). Analysis of unchallenged animals did not detect any podocin protein within several urine samples. Overall, ∼40% of *Ildr2^PodKO^* animals per challenge had detectable podocyte proteins in their urine, compared to only one of the *Ildr2^Pod/+^* animals (Fig. 4D). Cleaved caspase-3 analysis showed no significant increase in podocyte cell death at 24 hours post challenge (Fig. S4). Lastly, we assessed whether the challenge and podocyte loss induced any significant changes to kidney function. Even with the stress on kidney function and increase in podocyte detachment in *Ildr2^PodKO^* mice, we surprisingly did not find any significant differences in either proteinuria or plasma creatinine levels between groups (Fig. S5).

**Figure 4.**
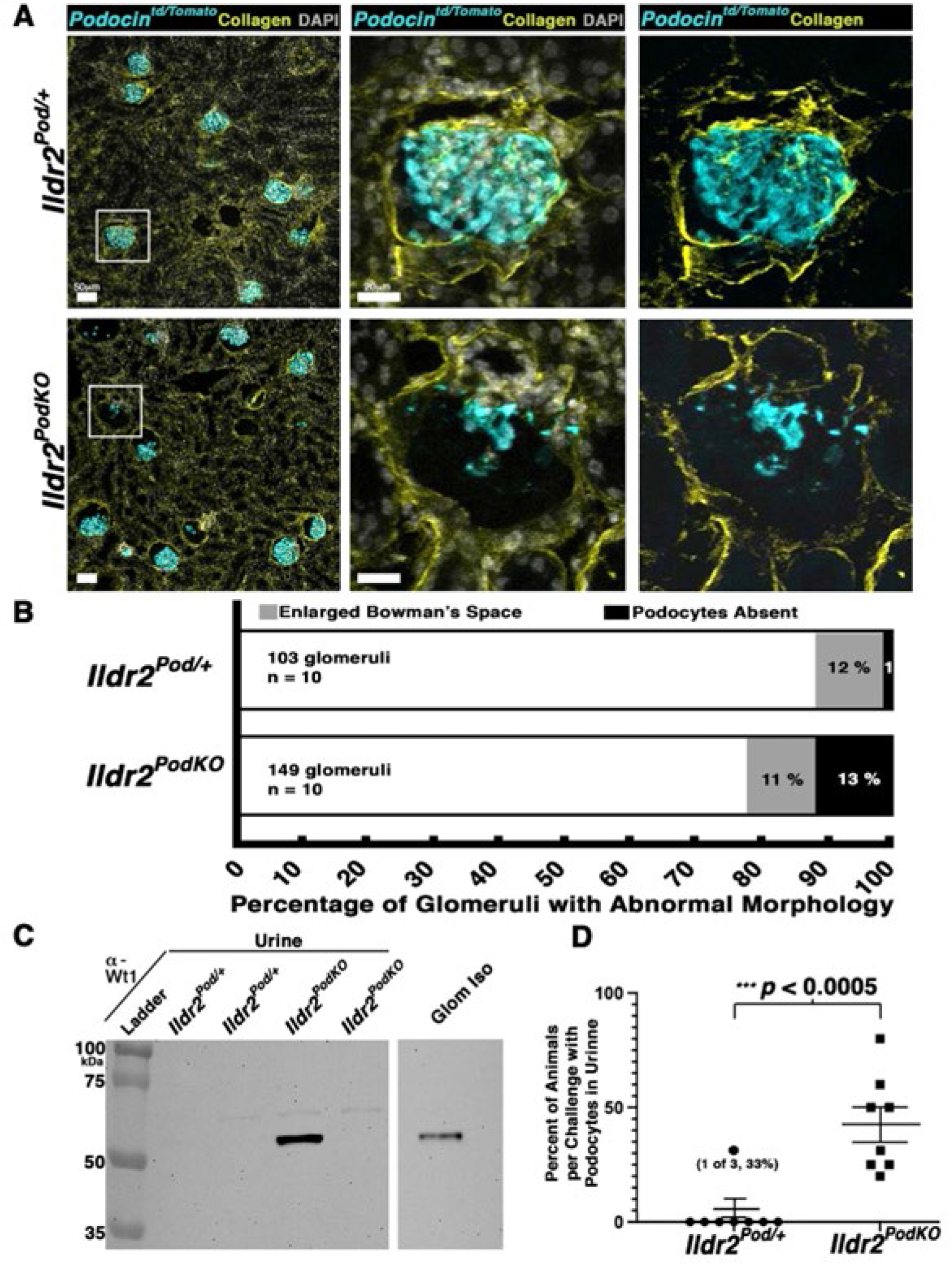
Challenge to *Ildr2^PodKO^* kidney function results in podocyte loss and detection in the urine. A) Immunohistological analyses of mice challenged with 10% body weight saline show podocyte loss from a subset of glomeruli. Podocytes are labeled with the tdTomato reporter crossed onto the *Ildr2^PodKO^* and *Ildr2^Pod/+^*backgrounds and is driven by *Podocin-Cre^tg^* (*Podocin^tdTomato^*). Boxed areas are shown in the adjacent zoomed images. Collagen IV (yellow), Tomato+ podocytes (cyan), and DAPI (grey) are labeled. Scale bar left=50 µm; scale bar in zoomed images=20 µm. B) Quantification of glomeruli with abnormal morphology after challenge shows a significant increase in the percentage of *Ildr2^PodKO^* glomeruli missing podocytes. C) Western blot of urine shows Wt1 signal in a sample from a challenged *Ildr2^PodKO^* animal, suggesting loss of podocytes into the urine. Isolated glomeruli (Glom Iso) are used as a control. D) Plot of the percentage of animals with podocyte proteins in their urine showing a significant percentage of *ldr2^PodKO^* animals have detectable podocyte proteins.

## Discussion

In this study we set out to determine the role of Ildr2 in podocyte function. We identified ultrastructural changes to podocyte architecture and thickening of the GBM in the absence of *Ildr2*. Additionally, we identified histological changes that suggest excess collagen accumulation occurs in *Ildr2^PodKO^* kidneys. Despite these phenotypes, podocyte junctions and their integrity are maintained in these animals. Stress from a challenge to kidney function induces podocyte loss in nearly half of these animals, however kidney function is still preserved. Altogether, we have identified a role for Ildr2 in maintaining proper podocyte architecture although it is not strictly required for the maintenance of barrier integrity.

Ildr2 is a component of tTJs and recent studies have shown that Ildr2/ILDR2 protein localization is altered in aging and disease^15,21^. These data suggest Ildr2/ILDR2 may be important to the maintenance of glomerular barrier integrity, and that these rearrangements into a more bicellular junctional pattern is an attempt to prevent further disease progression. Alternatively, Ildr2/ILDR2 redistribution may be a sign of disease progression and make the podocytes more vulnerable to further injury and loss, and lead to physiological changes such as proteinuria. Further analyses of aged and diseased tissues that lack Ildr2 will be important to help sort out its role in these processes.

The absence of any overt proteinuria or altered physiology in our *Ildr2^PodKO^* animals was surprising due to the effacement and histological changes we observed. However, if the observed phenotypes are not representative of the majority of glomeruli then kidney function may still be preserved. Exhaustive TEM and histological analyses would be necessary to assess whether this is the case. There may also be compensatory mechanisms that we were not able to uncover in our analyses. The resilience of the animals to functional challenge may suggest this is the case.

In depth gene expression analyses could provide additional clues into these potential players. On the other hand, tTJs that contain Ildr2 may not be essential to maintaining glomerular barrier integrity. With slit diaphragm and other cellular junctions largely still intact, the podocytes could still retain sufficient integrity to maintain function and buffer against any stresses. Although we do observe some podocyte loss in *Ildr2^PodKO^* animals with challenge, suggesting they may be sensitized, a more significant insult is likely necessary to tip the scales and lead to altered function. To the best of our knowledge, there is one other description of “extra” electron dense strands or filaments, aside from the slit diaphragm, protruding from the foot process ^11^ . In this study, they denote that foot processes can have numerous (>2) “punctate filamentous protrusions”. Other studies have noted “cross bridges” or “cross strands”^5,31^. In the *Ildr2^PodKO^* animals these protrusions may represent remnants of failed protein-bridges. Alternatively, these electron dense strands may be an entirely separate junctional complex aside from the slit diaphragm that represent an attempt to preserve the barrier in the absence of Ildr2 and proper tTJs. Further investigation is necessary to identify their composition and functional significance, if any.

While the exact role of tTJs in podocytes remains elusive, our studies have begun to shed light onto the full repertoire of junctional components that help maintain podocyte architecture. We speculate that the loss of or alterations to Ildr2 function may further sensitize already stressed podocytes in disease and aging and could represent a therapeutic target. Further investigations are necessary to better delineate the role of Ildr2 in kidney homeostasis, injury, and disease.

## Methods

### Animals

All animal use and husbandry procedures were performed according to the guidelines of the National Institutes of Health Guide for the Care and Use of Laboratory Animals and were approved by the Institutional Animal Care and Use Committees of The University of North Carolina at Chapel Hill (protocol 22-136). Mice were maintained in standard housing with corncob bedding, paper square nestlets, and hut enrichment on a 12-hour light–dark cycle at a room temperature of ∼22 °C. Mice had ad libitum access to food and water. *Podocin-Cre^tg^* (JAX stock #:008205) and *R26^tdTomato^* (JAX stock #:007914) mice were obtained from the Jackson Labs^24^ ^32^. *Ildr2^fl/fl^ mice* were a generous gift from Dr. Rudolph Leibel^23^. *R26^tdTomato^* was crossed in as necessary (denoted in text where included) to enable podocyte visualization and detection. All mice were maintained on a C57Bl6/J (JAX stock #:000664) background. No sex differences were identified in phenotypes upon initial assessments of males and females; males were predominantly utilized due to their availability. Mice were 3-5 months of age unless otherwise noted.

### Genotyping

Mouse tail and/or ear clip(s) were taken and dissociated with Viagen DirectPCR Lysis Reagent containing 10µg/mL proteinase K incubated at 55°C overnight and subsequently denatured at 95°C for 10 minutes. PCR was run with (Tannealing = 63.5°C), elongation for 40 seconds, for 35 cycles. Primer sequences utilized for genotyping are provided in Supplemental Table 1.

### Transmission Electron Microscopy (TEM)

The outer cortex of 5 month-old male littermate mice, *Ildr2^Pod/+^* and *Ildr2^PodKO^*, were isolated and protocols for TEM preparation and imaging were similar to that performed previously^15^. Briefly, kidneys were fixed for 1 hour at room temperature in 4% paraformaldehyde (PFA) in 0.15M sodium phosphate buffer pH 7.4. Samples were then washed 3 x 15 minutes (min) in 0.1M cacodylate buffer, stained in 1% OsO_4_ for 1 hour, followed by 3 x 15 min washes in ddiH20 and one wash in 50% ethanol for 5 min. Then samples were en-bloc stained with 2% uranyl acetate for 30 min. Successive dehydration washes were performed in 50, 70 and 95% ethanol 2x each for 10 min. Samples were then transferred to Wheaton glass jars where propylene oxide washes were performed 2 x 15 min each. A solution of 50/50 propylene oxide/EPON was applied overnight. The following day 25/75 propylene oxide/EPON was applied for 2 hours. Then 100% EPON was applied for 2 hours and fresh 100% EPON was applied for at least 24 hours and allowed to completely polymerize at 60°C. Blocks were trimmed to the tissue and 500µm sections were cut using a Leica EM UC7 (Leica Microsystems) and a Diatome diamond knife (Electron Microscopy Sciences). Sections were mounted on glass slides, stained with 1% toluidine blue O in 1% sodium borate, and regions with glomeruli were selected and trimmed. Ultrathin sections (80nm) were cut and mounted on 300 mesh copper grids. The grids were stained with 2% ethanolic uranyl acetate for 8 min, followed by staining with lead citrate for 5 min to add additional contrast. Tissue samples were visualized using a Philips Tecnai 12 at 120 kV and images were taken using a Gatan Rio 16 CMOS camera with Gatan Microscopy Suite software. Sample prep and imaging was performed at the UNC Hooker Imaging Core.

### Glomerular basement membrane (GBM) thickness measurements

Image J was utilized to measure from the bottom of the foot process atop the GBM, i.e. from the lamina rara externa layer, spanning the lamina densa, to the bottom of the lamina rara interna layer atop the endothelium. A blinded analysis was performed and at least three separate foot processes per podocyte were measured then averaged for a single GBM thickness measurement. A single data point represents the average thickness of the GBM in one podocyte (*Ildr2^Pod/+^* n=12, *Ildr2^PodKO^* n=9). A two-way t-test was performed to identify significant differences between control and *Ildr2^PodKO^* podocytes.

### Immunofluorescence

Kidneys were dissected in cold filter-sterilized PBS, cut longitudinally, and fixed at 4°C on a rocking platform for 45min-1hr in 4% paraformaldehyde (PFA) in PBS. Samples were washed twice in PBS, placed in 30% sucrose/PBS overnight, and embedded in Optimal Cutting Temperature (OCT) Embedding Medium (Fisher). Kidneys were sectioned at 12μm or 30μm (Fig. S4A) on a Leica Cryostat CM 1850. Tissue sections were blocked in 3% donkey serum, 1% bovine serum albumin (BSA), 1x PBS, with 0.1% TritonX-100 for 90 min at room temperature. Tissue sections were incubated in primary antibodies diluted in blocking buffer for 3-4 hours. Primary antibodies utilized include: rabbit anti-podocin (Invitrogen, PA5-79757) [1:500], goat anti-Collagen IV (Millipore, AB769) [1:1000], goat anti-mNephrin (R&DSystems AF3159) [1:50], rabbit anti-Cleaved Caspase-3 (CC3) (Cell Signaling, 9661S) [1:500], rabbit anti-ZO-1 (Proteintech, 21773-1-AP) [1:500], and mouse anti-Vinculin (Novus Biologicals, NB600-1293) [1:500]. Slides were then rinsed 3 x 5 min in 1x PBS + 0.1% TritonX-100 (PBST), after which they were incubated with the respective Alexa Fluor conjugated secondary antibodies (488, 568, or 647; Life Technologies) diluted (1:1000) in blocking buffer for 1 hour at room temperature. The tissue sections were then rinsed 2 x 5 min with 1x PBST, 1x 3–5 min with PBS + 1ng/mL DAPI and mounted in ProLong Gold antifade reagent (Life Technologies).

Images were acquired on a Zeiss 880 confocal microscope equipped with Airyscan super-resolution and spectral imaging on Zen Microscopy Suite version 2.3 Sp1 that is part of the UNC Hooker Imaging Core. Z-stack images for Fig. S4A were acquired in 5µm steps over 30µm to ensure that sectioning artifacts were not contributing to the absence of podocytes in our *Ildr2^PodKO^* challenged animals.

### Periodic Acid Shift (PAS) histology

Kidneys of 7-month-old male *Ildr2^Pod/+^* and *Ildr2^PodKO^* mice were dissected in cold filter-sterilized PBS and fixed at 4°C on a rocking platform in 4% PFA/PBS for 1 hour. Kidneys were washed with cold PBS and processed for paraffin embedding using standard protocols by the UNC Histology Research Core Facility. Kidney sections from each genotype, at 3µm thickness, were adhered on the same slide to control for staining variability. The tissue was deparaffinized with xylene then subjected to ethanol rehydration in successive washes with 100%, 95%, 70% and finally in water with two washes of each solution for 3-5 min per wash. Slides were then immersed in Periodic Acid Solution (Abcam) for 10 min and rinsed in distilled water. Tissue sections were then immersed in Schiff’s solution for 30 min and rinsed in hot running water. Kidney sections were then rinsed in distilled water at room temperature. Slides were stained in Hematoxylin for 3 min. Slides were rinsed in running tap water for 3 min, followed by one rinse in distilled water. Tissue sections were then successively dehydrated in ethanol solutions, and mounted in Cytoseal XYL (Epredia), and imaged on a Leica DMi8.

### Sirius Red histology

*Ildr2^Pod/+^* and *Ildr2^PodKO^* kidney sections from 7-month-old male mice were deparaffinized and hydrated similar to PAS histology procedures with control and mutant tissue adhered on the same slide. Tissue sections were Incubate in 0.2% phosphomolybdic acid for 2 min. Kidney tissue was then rinsed 2 x 2 min in ddH2O. A solution of 0.1% Picrosirius Red (Fisher) in saturated aqueous picric acid was applied for 30 min at room temperature. Tissue samples were washed 2 x 5 min in 100% ethanol, mounted, and imaged on a Leica DMi8.

### Albuminuria measurements

*Ildr2^PodKO^* and littermate control *Ildr2^Pod/+^*mice at 3-4 months of age were placed in metabolic cages for 24 hours with ad libitum access to food and water. Urine was collected for the 24-hour period and assayed for total protein concentration utilizing a standard Bradford assay.

### Saline challenge

*Ildr2^PodKO^* and littermate control *Ildr2^Pod/+^* mice were subcutaneously injected with a volume of 10% body weight, sterile saline solution (0.9% NaCl, prewarmed). The needle was slowly removed to prevent leakage, and excess solution wiped away. The mice were then immediately placed in metabolic cages for 24 hours with ad libitum access to food and water. Urine was collected for the duration of this 24-hour period for subsequent analyses.

### Creatinine measurements

Blood plasma was collected 24 hours after saline challenge. Serum was spun down at 10,000g for 10 min and plasma was removed. Aliquots of 30µL of plasma were plated in duplicate into a 96-well round bottom plate. The BioAssays System Creatinine (QuantiChrom) reagents were mixed in equal parts and added to each well rapidly with a multichannel pipette. The plate was immediately read at 510 nm and then read again 5 min later. The Jaffe method was utilized to determine the relative creatinine concentration of each sample and as noted in Keppler et al., 2007^33^, this method of analysis lends to higher readings.

### Western blot of urine

Urine collected for the 24-hour period immediately following challenge was measured, centrifuged, and a 30µL urine aliquot near the bottom of the tube was collected for analysis. Glomeruli preps as described in Gerlach et al., 2023^15^ were used as control tissue. Samples were denatured in 4x Lamelli buffer with heating at 95°C for 20 minutes and loaded into a 10% Tris-Glycine Novex WedgeWell gel (Life Technologies) run in 25mM Tris-HCl, 190mM Glycine, 0.1% SDS, pH 8.3. Proteins were transferred from gel to nitrocellulose membrane in 25mM Tris-HCl, 190mM Glycine, and 20% methanol. The membrane was blocked in 3% bovine serum albumin in 1x TBST (25mM Tris-HCl pH 7.5, 150mM NaCl, 0.1% Tween-20) for 1 hour at room temperature. Primary antibodies were diluted in 3% BSA/TBST and incubated at 4°C overnight with rocking/shaking. Primary antibodies utilized include: rabbit anti-Podocin (Invitrogen, PA5-79757) [1:500], chicken anti-RFP (Rockland, 600-901-379) [1:1000], and rabbit ant-Wilms Tumor (Wt1, Abcam) [1:1000]. The following day 3 x 15 minute washes were performed with TBST. The appropriate secondary antibodies conjugated to HRP were diluted 1:1000 in TBST and incubated with the membrane for 1 hour at room temperature, followed by 3 x 15 minute washes with TBST. Membranes were developed with enhanced chemiluminescence substrate (ECL) and visualized on an iBright FL1000 (Invitrogen).

### qPCR

Glomerular preps were performed as in Gerlach et al., 2023^15^. RNA extraction, cDNA synthesis, and qPCR were carried out similar to Honeycutt et al., 2023^34^.

## Acknowledgements

We would like to acknowledge the many people and core services from the University of North Carolina at Chapel Hill who helped with this investigation. We specifically would like to thank Paul Risteff for training in TEM, aid with image acquisition, and sample preparation and the Histology Core Services for histological sample preparation. We thank Dr. Rudolph Leibel (Columbia University) for the generous sharing of the *Ildr2^flox^* mice. We would further like to acknowledge and thank the members of the O’Brien lab for critical feedback on these studies.

## Author Contributions

LLO and GFG conceptualized and designed the study. GFG performed the experiments, interpreted and analyzed the data, and wrote the manuscript. YX performed blinded GBM thickness measurements. LLO supervised research, helped design and guided experiments, helped interpret data, and edited the manuscript.

## Supplemental Figures and Tables

**Supplemental Figure 1.**
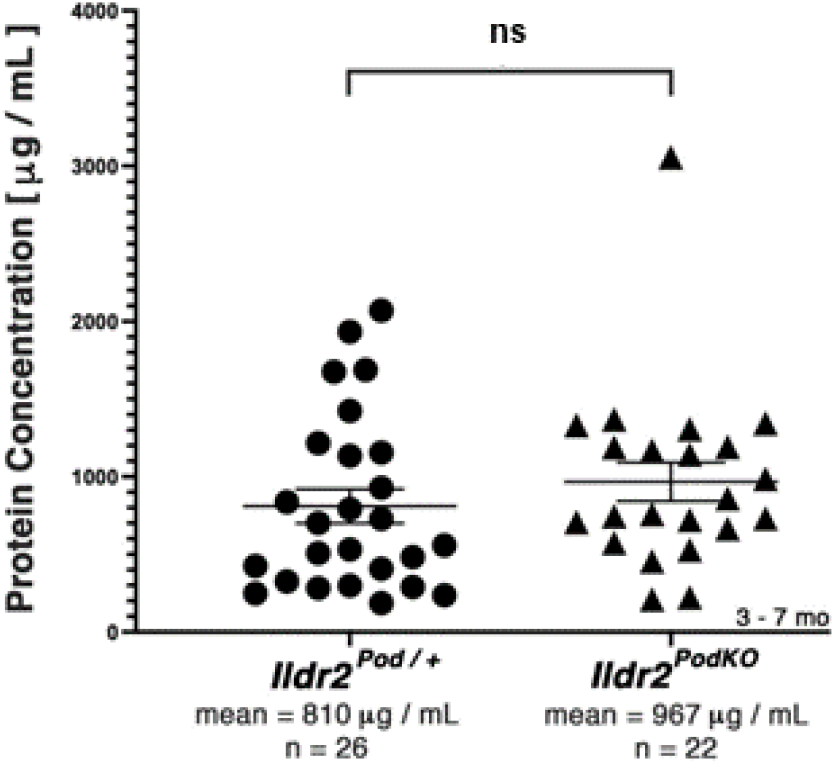
Expression of junctional components is not altered in *Ildr2^PodKO^* animals. *Ildr2^Pod/+^* and *Ildr2^PodKO^* mice were analyzed for gene expression differences via qPCR. No significant differences in the expression of *Tjp1* (ZO-1) nor *Nphs2* (podocin) were identified. ns=non-significant.

**Supplemental Figure 2.**
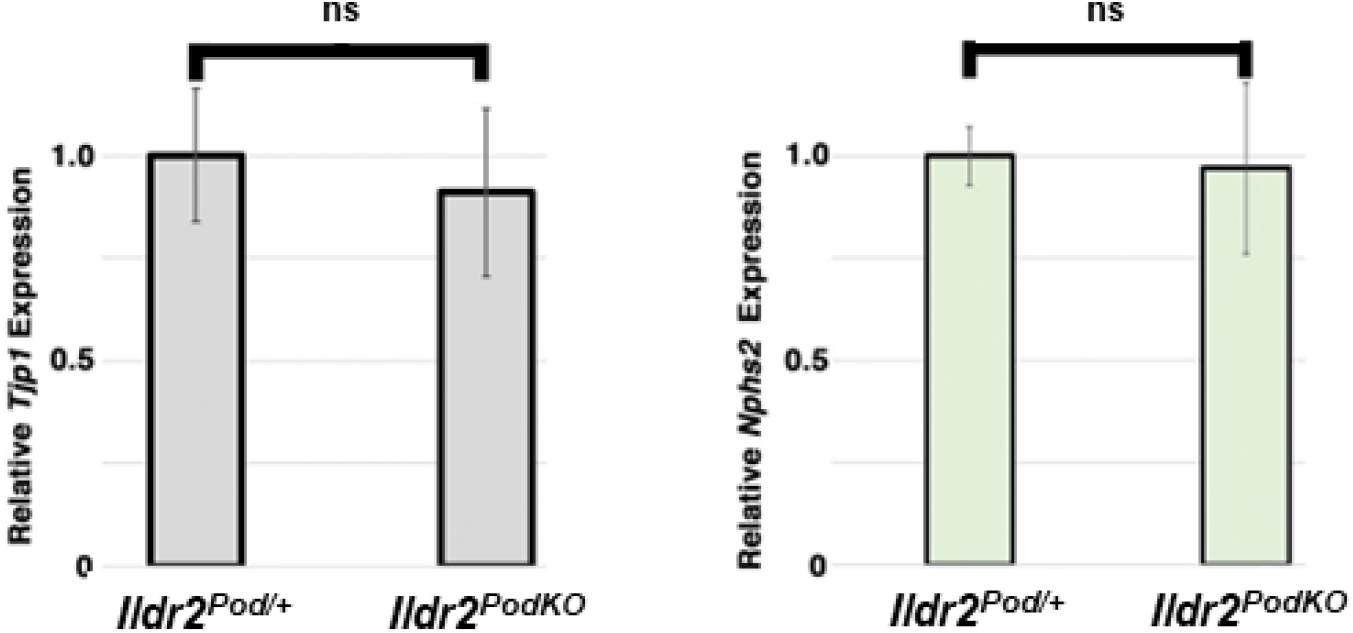
I*l*dr2PodKO animals do not exhibit proteinuria under normal physiological conditions. Urine collected from *Ildr2^Pod/+^*and *Ildr2^PodKO^* animals was assayed for total protein concentration utilizing a standard Bradford assay and no significant difference between control and *Ildr2^PodKO^* animals was detected. ns=non-significant.

**Supplemental Figure 3.**
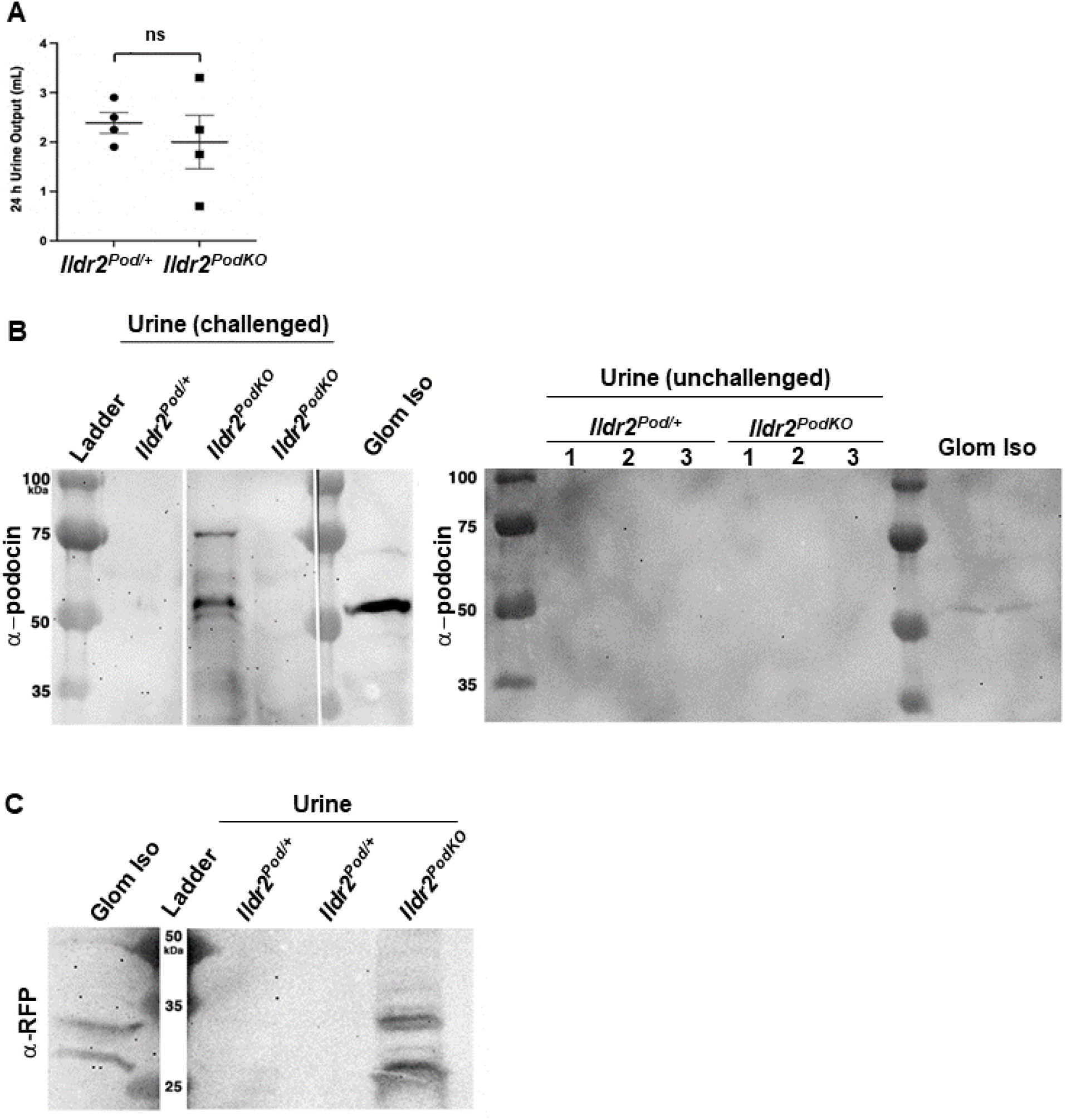
Challenge to kidney function leads to podocyte loss and detection in the urine of *Ildr2^PodKO^* animals. A) Urine volume measured after 24 hours of collection from mice injected with 10% body weight of a 0.9% saline solution showing no significant difference between *Ildr2^Pod/+^* and *Ildr2^PodKO^* animals. ns=non-significant. B) Left: Western blot of urine shows podocin signal in a sample from a challenged *Ildr2^PodKO^* animal, suggesting loss of podocytes into the urine. Isolated glomeruli (Glom Iso) are used as a control. Right: Western blot of urine from unchallenged animals shows lack of any significant podocin signal in samples from *Ildr2^PodKO^* mice. Isolated glomeruli (Glom Iso) are used as a control. C) Western blot of urine shows RFP/Tomato signal in a sample from a challenged *Ildr2^PodKO^;R26^tdTomato^*animal, suggesting loss of podocytes into the urine. Isolated glomeruli (Glom Iso) are used as a control.

**Supplemental Figure 4.**
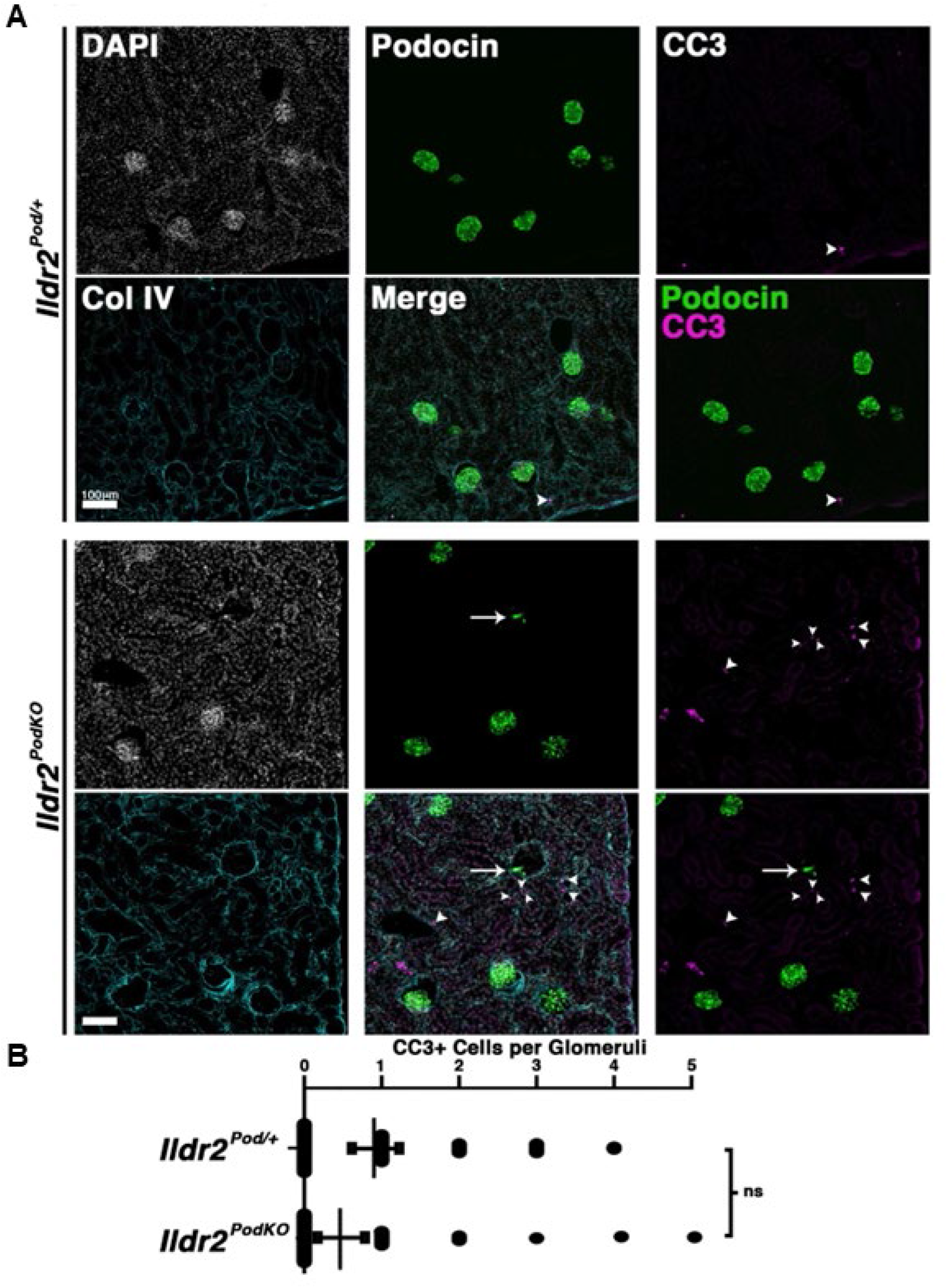
Challenge to kidney function induces podocyte loss in *Ildr2^PodKO^* animals without a detectable increase in apoptotic cell death in glomeruli. A) Cryosectioned kidneys immunostained for podocytes (podocin), apoptotic cells (CC3), and glomerular architecture (Collagen IV, ColIV) show normal glomerular organization in controls and examples of *Ildr2^PodKO^*glomeruli lacking podocytes (arrow) in challenged animals. Arrowheads point to apoptotic cells. In *Ildr2^PodKO^* animals, CC3+ cells are found outside glomeruli. Scale bar=100µm. B) Quantitation of CC3+ cells within glomeruli reveals no significant differences between *Ildr2^Pod/+^* and *Ildr2^PodKO^* animals. ns=non-significant.

**Supplemental Figure 5.**
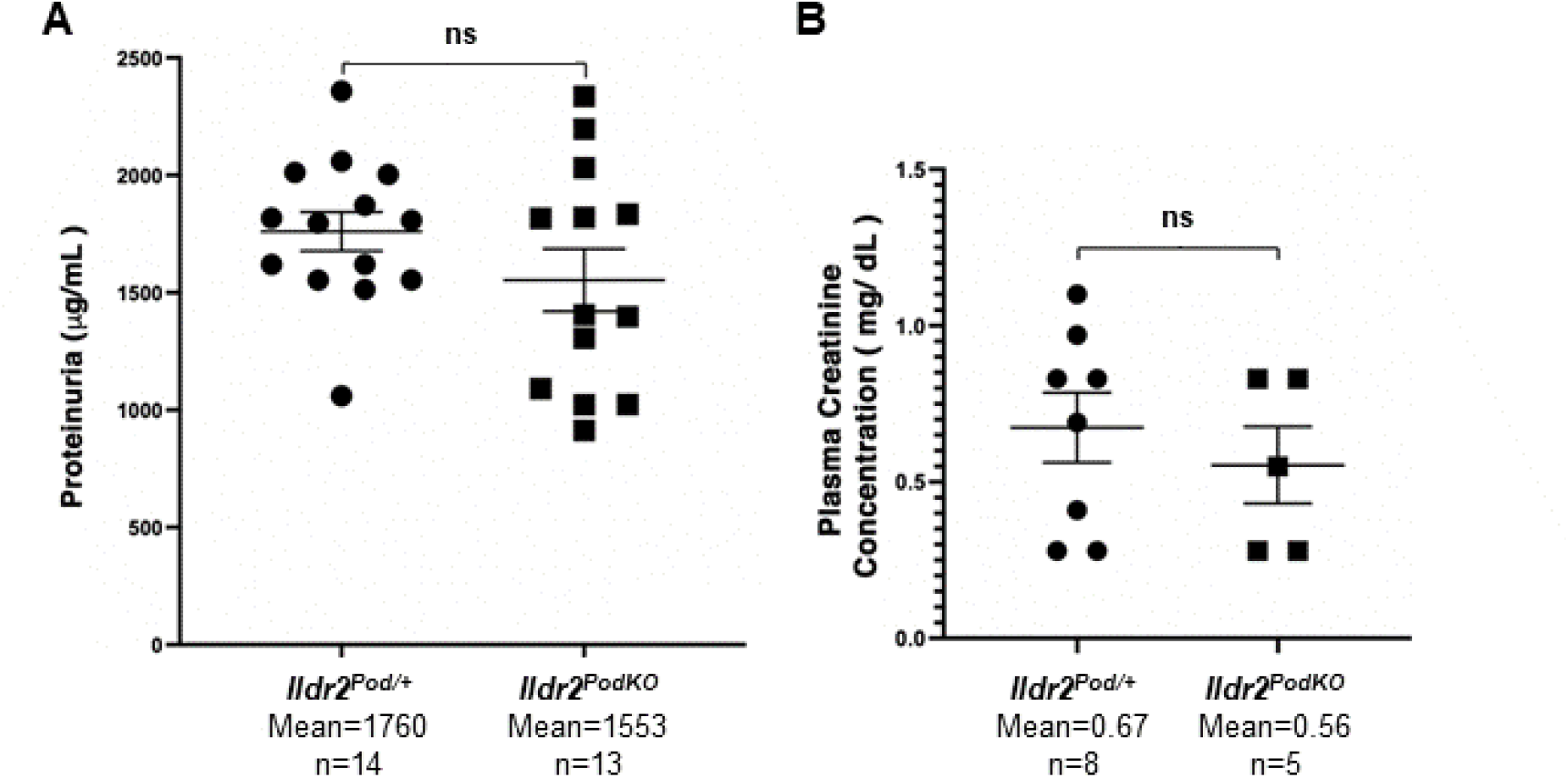
I*l*dr2PodKO animals do not exhibit physiological changes following challenge. A) Proteinuria analysis, utilized as a readout of glomerular function, was not significantly different between controls and *Ildr2^PodKO^* animals 24 hours post challenge. B) Serum creatinine measured 24 hours post challenge shows no significant differences between controls and *Ildr2^PodKO^* animals. ns=non-significant.

**Supplemental Table 1.**
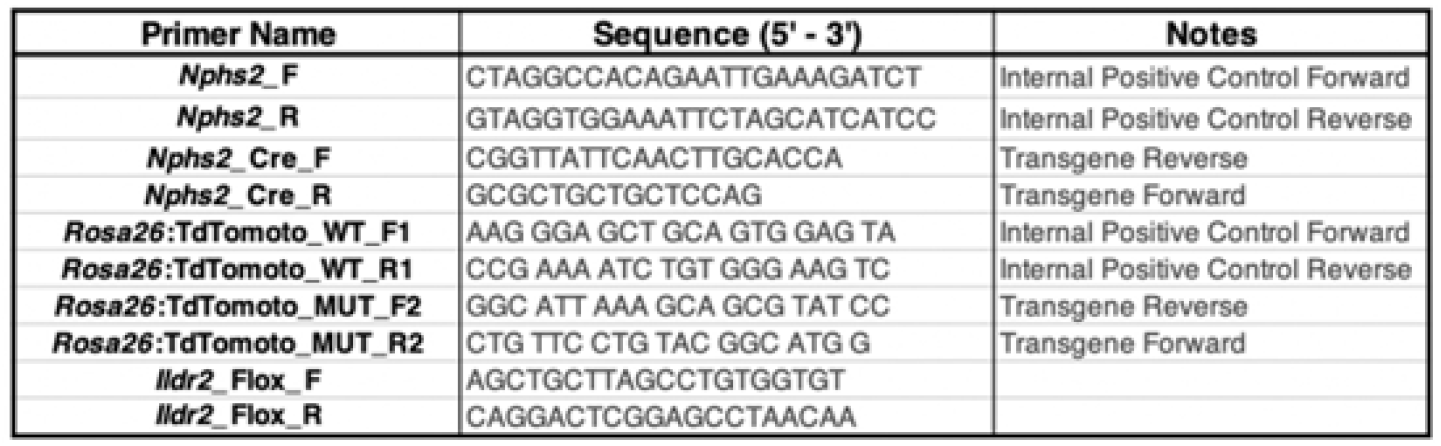
Genotyping primers utilized in this study.

