## Supplemental Figures and Tables for "The tricellular junctional protein Ildr2 helps maintain glomerular podocyte architecture"

### Supplemental Figure 1

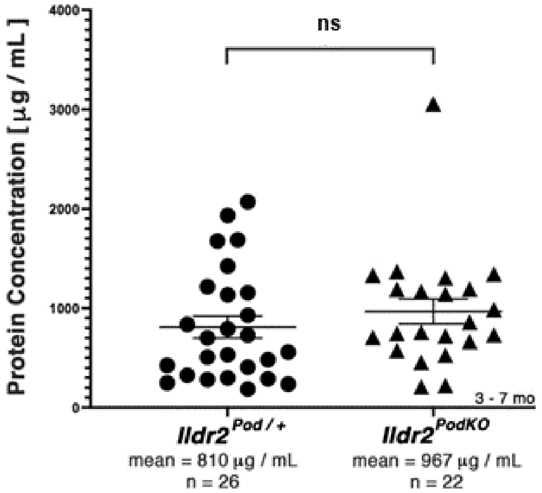


**Supplemental Figure 1. Expression of junctional components is not altered in *Ildr2*$^{PodKO}$ animals.** *Ildr2*$^{Pod/+}$ and *Ildr2*$^{PodKO}$ mice were analyzed for gene expression differences via qPCR. No significant differences in the expression of *Tjp1* (ZO-1) nor *Nphs2* (podocin) were identified. ns=non-significant.

### Supplemental Figure 2

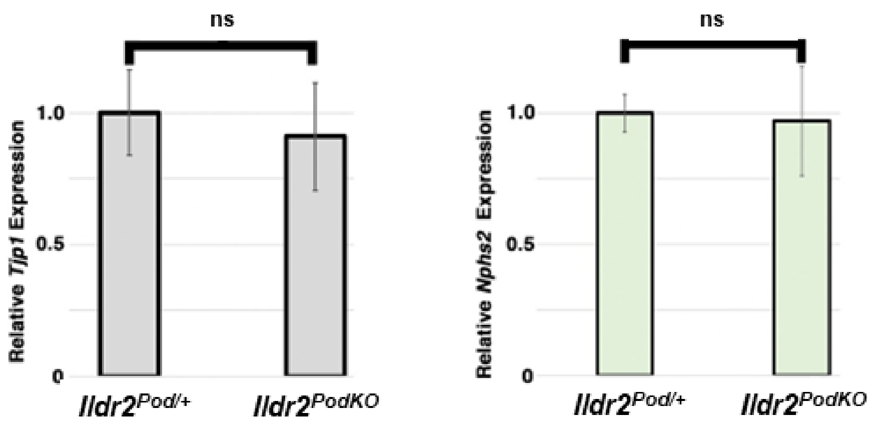


**Supplemental Figure 2. *Ildr2$^{PodKO}$* animals do not exhibit proteinuria under normal physiological conditions.** Urine collected from *Ildr2$^{Pod/+}$* and *Ildr2$^{PodKO}$* animals was assayed for total protein concentration utilizing a standard Bradford assay and no significant difference between control and *Ildr2$^{PodKO}$* animals was detected. ns=non-significant.

### Supplemental Figure 3

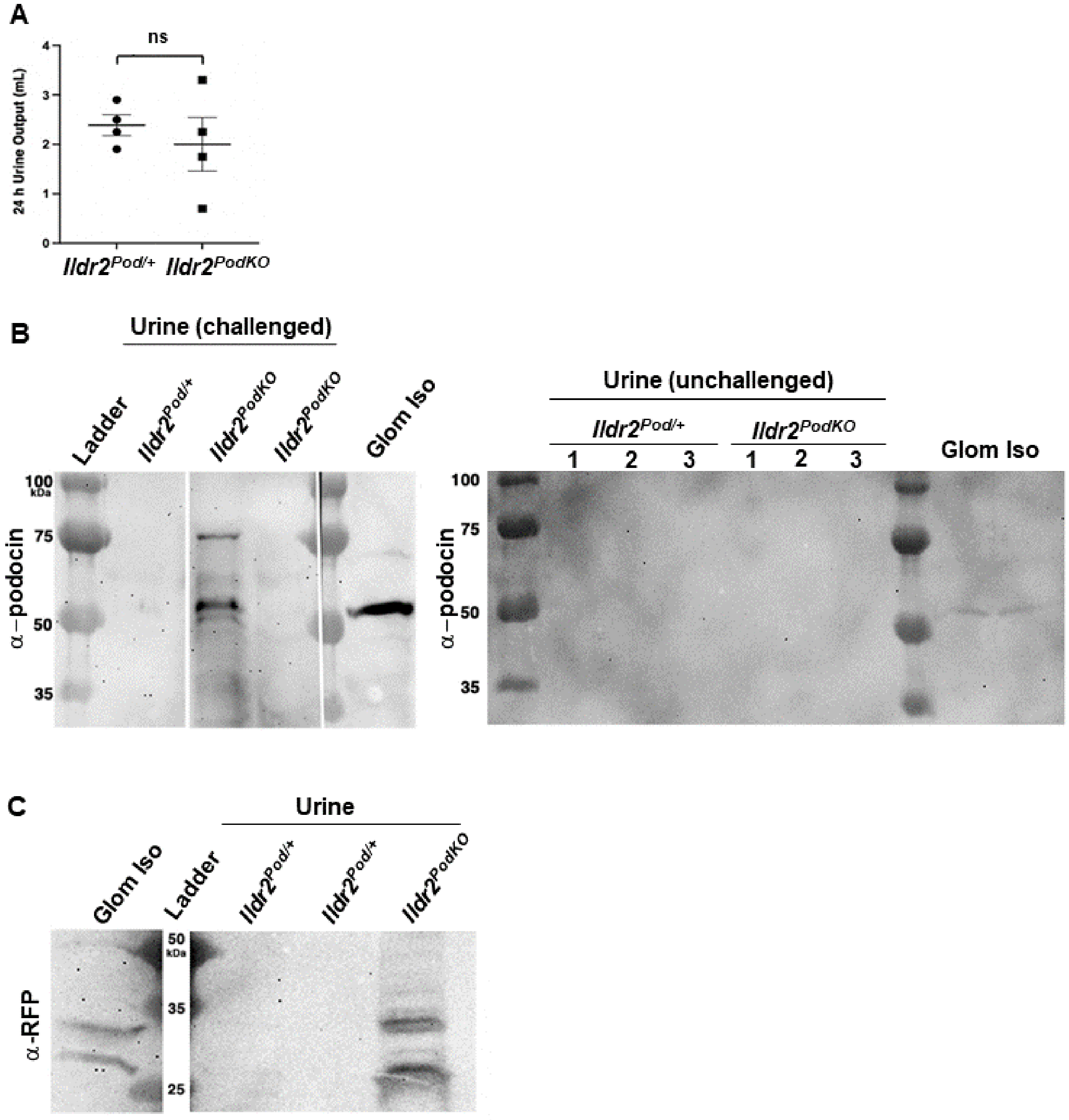


**Supplemental Figure 3. Challenge to kidney function leads to podocyte loss and detection in the urine of *Ildr2*$^{PodKO}$ animals.** A) Urine volume measured after 24 hours of collection from mice injected with 10% body weight of a 0.9% saline solution showing no significant difference between *Ildr2*$^{Pod/+}$ and *Ildr2*$^{PodKO}$ animals. ns=non-significant. B) Left: Western blot of urine shows podocin signal in a sample from a challenged *Ildr2*$^{PodKO}$ animal, suggesting loss of podocytes into the urine. Isolated glomeruli (Glom Iso) are used as a control. Right: Western blot of urine from unchallenged animals shows lack of any significant podocin signal in samples from *Ildr2*$^{PodKO}$ mice. Isolated glomeruli (Glom Iso) are used as a control. C) Western blot of urine shows RFP/Tomato signal in a sample from a challenged *Ildr2*$^{PodKO}$*;R26*$^{tdTomato}$ animal, suggesting loss of podocytes into the urine. Isolated glomeruli (Glom Iso) are used as a control.

### Supplemental Figure 4

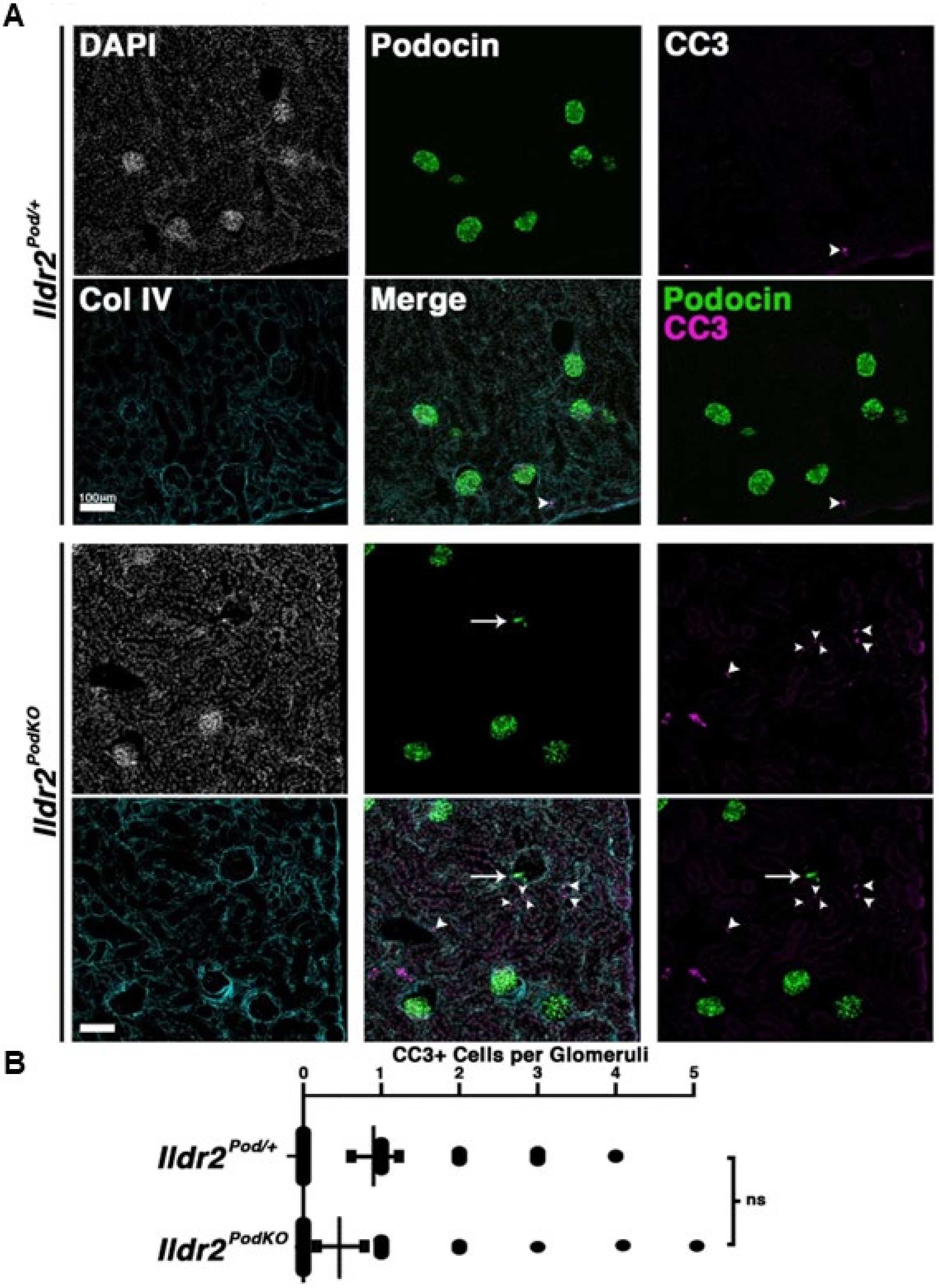

**Supplemental Figure 4. Challenge to kidney function induces podocyte loss in *Ildr2*$^{PodKO}$ animals without a detectable increase in apoptotic cell death in glomeruli.** A) Cryosectioned kidneys immunostained for podocytes (podocin), apoptotic cells (CC3), and glomerular architecture (Collagen IV, ColIV) show normal glomerular organization in controls and examples of *Ildr2*$^{PodKO}$ glomeruli lacking podocytes (arrow) in challenged animals. Arrowheads point to apoptotic cells. In *Ildr2*$^{PodKO}$ animals, CC3+ cells are found outside glomeruli. Scale bar=100μm. B) Quantitation of CC3+ cells within glomeruli reveals no significant differences between *Ildr2*$^{Pod/+}$ and *Ildr2*$^{PodKO}$ animals. ns=non-significant.

### Supplemental Figure 5

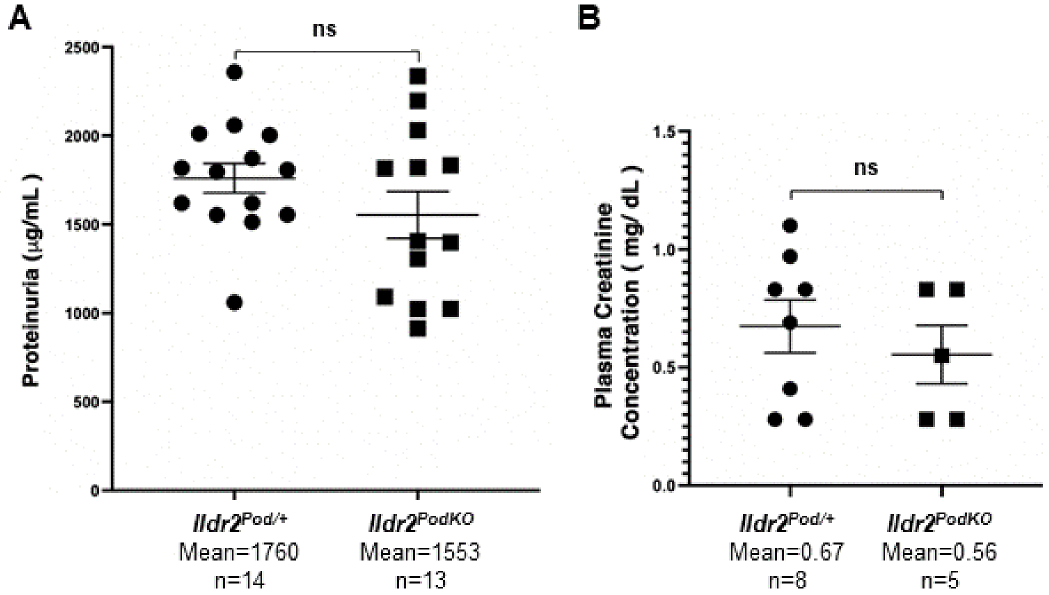


**Supplemental Figure 5. *Ildr2$^{PodKO}$* animals do not exhibit physiological changes following challenge.** A) Proteinuria analysis, utilized as a readout of glomerular function, was not significantly different between controls and *Ildr2$^{PodKO}$* animals 24 hours post challenge. B) Serum creatinine measured 24 hours post challenge shows no significant differences between controls and *Ildr2$^{PodKO}$* animals. ns=non-significant.

### Supplemental Table 1

| Primer Name | Sequence (5' - 3') | Notes |
|---|---|---|
| *Nphs2*_F | CTAGGCCACAGAATTGAAAGATCT | Internal Positive Control Forward |
| *Nphs2*_R | GTAGGTGGAAATTCTAGCATCATCC | Internal Positive Control Reverse |
| *Nphs2*_Cre_F | CGGTTATTCAACTTGCACCA | Transgene Reverse |
| *Nphs2*_Cre_R | GCGCTGCTGCTCCAG | Transgene Forward |
| *Rosa26*:TdTomoto_WT_F1 | AAG GGA GCT GCA GTG GAG TA | Internal Positive Control Forward |
| *Rosa26*:TdTomoto_WT_R1 | CCG AAA ATC TGT GGG AAG TC | Internal Positive Control Reverse |
| *Rosa26*:TdTomoto_MUT_F2 | GGC ATT AAA GCA GCG TAT CC | Transgene Reverse |
| *Rosa26*:TdTomoto_MUT_R2 | CTG TTC CTG TAC GGC ATG G | Transgene Forward |
| *Ildr2*_Flox_F | AGCTGCTTAGCCTGTGGTGT | |
| *Ildr2*_Flox_R | CAGGACTCGGAGCCTAACAA | |

**Supplemental Table 1. Genotyping primers utilized in this study.**
